# Cortical Serotonin Type 2A Receptor Activation Shields Episodic-like Memories from Retroactive Interference in Rodents

**DOI:** 10.1101/2025.06.25.661375

**Authors:** JF Morici, Mariana Imperatori, Pablo Koss, MB Zanoni, A Sacson, P Bekinschtein, NV Weisstaub

**Affiliations:** Instituto de Neurociencia Cognitiva y Traslacional (INCYT), CONICET, Fundación INECO, Universidad Favaloro, Buenos Aires, Argentina; Sorbonne Université, CNRS, Inserm, Institut de Biologie Paris-Seine, Center of Neuroscience NeuroSU, F-75005 Paris, France; Instituto Tecnológico de Buenos Aires (ITBA), Buenos Aires, Argentina; Departamento de Fisiología, Biología Molecular y Celular “Dr. Héctor Maldonado” (FBMC), Universidad de Buenos Aires (UBA), Facultad de Ciencias Exactas y Naturales, Buenos Aires, Argentina

## Abstract

The acquisition of temporally proximate information can impair the brain’s ability to consolidate earlier experiences, resulting in retroactive interference (RI). Recognition-based behavioral paradigms are well-suited for investigating RI in rodents, particularly those involving sequential learning episodes. The medial prefrontal cortex (mPFC) integrates multimodal information relevant to the regulation of memory interference and is strongly modulated by the serotonergic system. Serotonin 2A receptors (5-HT2AR), which are densely expressed in the mPFC, have been shown to influence the retrieval of competing object-recognition memories. However, their role in other phases of memory processing, particularly in modulating RI, remains unclear. Using a modified version of the object recognition task designed to induce RI, combined with pharmacological manipulation of 5-HT2AR, we demonstrate that RI specifically impairs the object-related component of memory. Moreover, serotonin signaling through 5-HT2AR is necessary to prevent RI. Strikingly, activating 5-HT2AR before retrieval can rescue the expression of memories affected by RI, suggesting that RI may not erase memory traces but rather hinder their access.

## Introduction

The acquisition of temporally proximate information can impair the brain’s ability to form strong memories. During learning, changes in specific neurons and synapses give rise to a “neural trace” of the memory, which is distributed across brain areas depending on the features of the experience. When similar experiences occur close in time, they may recruit overlapping populations of neurons, resulting in *retroactive interference* (RI), a phenomenon that compromises the stability or accessibility of the earlier memory ^1,2^. Recognition-based behavioral tasks are well-suited for studying RI in rodents, as they allow investigation of competing, emotionally neutral memories. Using such paradigms, introducing a new object-context association, or even a novel object alone, during the consolidation phase of a previously acquired memory can induce RI ^3,4^. Interestingly, RI appears to depend on competition among original memory engrams (Autore et al., 2023) and engages multiple brain structures ^3–5^.

The medial prefrontal cortex (mPFC) integrates multimodal information and plays a key role in memory selection, during both encoding and retrieval ^6^. Evidence from animal and human studies indicates that the mPFC contributes to reducing memory interference during acquisition ^3,4,7^. The serotonergic system, a major neuromodulator of mPFC function, plays a central role in this process. In particular, the 5-HT2A receptor (5-HT2AR) is highly expressed in the mPFC and has been linked to mood regulation and cognitive functions ^8^. Previous studies have shown that mPFC 5-HT2AR activity is required to regulate the expression of competing object-recognition memories through top-down control of other brain regions during retrieval ^9,10^. However, whether the mPFC 5-HT2AR also plays a role in controlling memory interference during other phases of memory processing remains unclear. Although emerging evidence implicates 5-HT2AR in memory consolidation across species ^11–13^, its involvement specifically during retroactive interference in the consolidation phase has yet to be investigated.

To investigate the role of serotonergic modulation, particularly via the 5-HT2A receptor, in controlling interfering memories, we implemented an object recognition paradigm that induces retroactive interference. Using this approach, we manipulated 5-HT2AR activity in the mPFC to either prevent or promote RI-induced forgetting. Interestingly, we found that this forgetting affected not only the integrity of the object-context association but also the object memory itself. Moreover, activation of 5-HT2AR after the second learning event prevented RI-induced forgetting of the first object-context association.

## Materials and Methods

### Animals

A total of 167 male Wistar rats were used. At the time of the experiments, rats weighed between 200 and 250 g and were grouped. The animals were kept with water and food ad libitum under a 12 h light/dark cycle (lights on at 7:00 A.M.) at a constant temperature (22–23 °C. Experiments took place during the light phase of the cycle (between 10:00 A.M. and 5:00 P.M.) in quiet rooms with dim lighting. All experimental procedures were performed according to institutional regulations and following those of the National Animal Care and Use Committee of the Favaloro University (CICUAL Presentation DCT0204-16 / PICT 2015-0110)

### Behavioral experiments

#### Apparatus

We used two different-shaped mazes made of white acrylic. The first apparatus was a 50 cm wide X 50 cm long X 39 cm high rectangular maze. The second apparatus was a 60 cm wide X 40 cm long X 50 cm high rectangular maze. Each wall has a visual distinctive cue that is different from the other cues presented.

All contexts were designed to have identical surface areas to eliminate any differences due to arena size. Duplicate plastic, glass, and aluminum objects were used, thoroughly cleaned between phases, and randomly assigned to different phases of the experiments. The objects were securely attached to the apparatus floor with an odorless, reusable adhesive to prevent movement during each session. They were consistently positioned along the centerline of the maze, away from the walls, and equidistantly spaced. To the best of our knowledge, the objects had no natural meaning to the animals, as they were never associated with any form of reinforcement. The objects, floor, and walls were cleaned with 50% ethanol between trials.

#### Object-in-context

This three-trial task evaluates the congruency between an object and its context ^10,14,15^. In the training phase, animals are exposed to two identical object pairs, each presented in a different context. During the test, conducted 24 hours later, new copies of the objects are presented in pseudorandomly assigned contexts, resulting in one contextually “congruent” and one “incongruent” object. During habituation sessions, rats were allowed to explore each context for 10 min. In the first training session, two identical objects (A1 and A2) were placed in one of the arenas (context 1). The animals were introduced into the context facing the wall and allowed to explore for either 3 or 5 minutes, depending on whether they were undergoing short or long training. After this session, the animals were returned to their home cage. Following a 1-hour delay, they were placed in a second context with a new pair of objects (B1 and B2) for the same duration (3 or 5 minutes). The arenas were pseudorandomly assigned as context 1 or 2. During the test session, animals were reintroduced to context 1 or context 2 and were allowed to explore for 3 min one copy of object A and one copy of object B. This session occurred 3 hours after the training session for short-term memory tests, or 24 hours later for long-term memory tests. To facilitate comprehension, a schematic representation of each experiment is placed above the quantifications.

#### Behavioral analysis

During each session, the time spent exploring each object was recorded. In both the training and test phases, total exploration time for each object copy was analyzed. For the test phase, the Discrimination Index (DI) was calculated as follows: (time spent on the incongruent object – time spent on the congruent object) / total exploration time in the session. Exploration was defined as the animal directing its nose toward the object within 2 cm or touching it with its nose, while behaviors like turning around or sitting on the object were excluded. All experiments were recorded using Samsung HMX-F80 cameras positioned above the arenas to capture the entire space. Offline analysis was conducted using manual stopwatches for all phases by a trained experimenter who was blind to the experimental conditions.

### Surgery and drug infusions

The animals were deeply anesthetized using ketamine (Holliday, 80 mg/kg, i.p.) and xylazine (Konig, 8 mg/kg, i.p.), then positioned in a stereotaxic frame (Stoelting). The skull was exposed and adjusted to align the bregma and lambda on the same horizontal plane. After drilling small burr holes, one, two, or three pairs of guide cannulae (22 g) were bilaterally implanted into the medial prefrontal cortex (mPFC, AP: +3.20 mm, LL: ± 0.75 mm, DV: -3.50 mm). All coordinates were calculated using the bregma as a reference. Following the surgery, and until the infusions, a dummy injector (30 G) was placed into the guide cannulae to prevent obstruction. To verify the cannula placement, 24 hours after the conclusion of the behavioral experiments, animals were infused with 1 ml of methylene blue through the dummy cannulae. Fifteen minutes later, they were deeply anesthetized and sacrificed. The infusion sites were then histologically localized using magnifying glasses. No animals were excluded due to cannula misplacement.

On the training day, following the second session, the dummy injector was removed and a 30 G injection cannula, extending 1 mm below the guide cannula, was inserted. This injection cannula was attached to a 10 μl Hamilton syringe. Each animal received a bilateral infusion of 1 μl ofl Hamilton syringe. Each animal received a bilateral infusion of 1 μl Hamilton syringe. Each animal received a bilateral infusion of 1 μl ofl of either the appropriate drug or its vehicle into the mPFC. The drugs infused were: a selective 5-HT2aR antagonist, MDL 11,939 (MDL, 300 ng/µl, Tocris Bioscience, Pittsburgh), a 5-HT2 agonist, DOI (±)-DOI hydrochloride (DOI, 10ng/µl, SIGMA-ALDRICH), and a selective 5-HT2AR agonist TCB-2 (TCB, 4 µg/µl, Tocris Bioscience, Pittsburgh) or vehicle (5% DMSO in saline).

### Statistical analyses

Statistical analyses were performed with GraphPad 10. Behavioral data were analyzed using a two-tailed unpaired Student’s t-test when two groups were compared. For comparisons between two groups, a two-tailed Student’s t-test was used. Two-way ANOVA followed by a Sidak post hoc test, as indicated in the figure legends, was used when three or more groups were involved. In all cases, p-values were considered to be statistically significant when p <0.05. All data are presented as scatter plots, including the mean ± s.e.m.

## Results

We trained rats using the object-in-context (OIC) paradigm, which requires animals to identify, during the test session, the object that is incongruent with the context in which it is presented, based on prior experience. This task is traditionally used to model interference during the retrieval of episodic-like memories in rodents (Bekinschtein et al., 2013; Morici et al., 2018). However, because the OIC paradigm involves the encoding of two similar experiences in close temporal proximity, it also provides a useful model for studying interference during memory *storage*.

To examine this possibility, we used a within-subjects design and trained rats with a short training session (ST) of the OIC paradigm. Each week, rats underwent a training session in which they were first exposed to objects A in context 1 for 3 minutes, followed, after a fixed delay of either 15, 60, or 300 minutes, by exposure to objects B in context 2 for another 3 minutes. The following day, memory for either the first (A-context 1) or second (B-context 2) object-context association was tested, with the tested association chosen pseudo-randomly across animals. One week later, the same procedure was repeated with new contexts and objects (C in context 3, D in context 4), using the same delay assigned during the previous week. This time, the association that had not been tested the week before was tested (see behavioral protocol in **Fig. 1a**).

**Figure 1:**
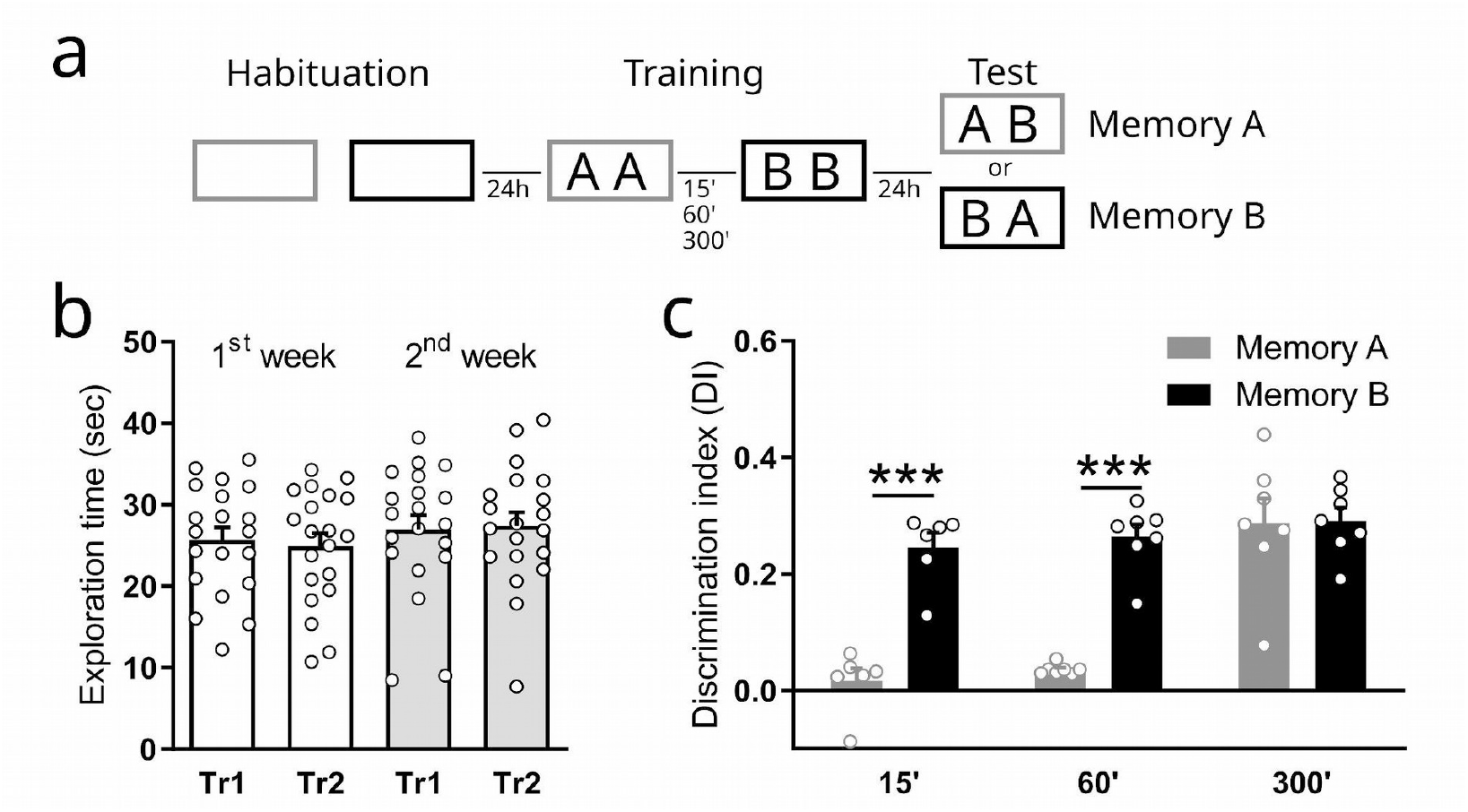
Object-in-context retroactive interference is a time-dependent process. Retroactive interference (RI) is a phenomenon observed when two learning experiences occur in a specific time frame. (a) Scheme of the object-in-context protocol. If an RI process occurs in this task, we should see a time-dependent amnesic effect over the first memory. (b) Exploration time for each object obtained during acquisition sessions. 2-way ANOVA, p_interaction_=0.26, F(1,19)=1.36. (c) Discrimination Index (DI) obtained during the test session. DI was calculated as (t_incongruent_ – t_congruent_)/(t_incongruent_ + t_congruent_). Both OIC memories were evaluated in all animals. Grey bars (Memory A) represent the memory of Tr1, while black bars (Memory B) represent the memory of Tr2. 2-way ANOVA followed by Sidak post hoc test. ****p_interaction_ < 0,0001, F(2,34)=13.12, ***p_post-hoc_ < 0,001, n=6-7 per group.

Notably, total object exploration times during training were comparable across sessions and weeks, indicating stable exploratory behavior (**Fig. 1b**). Memory performance was evaluated using a Discrimination Index (DI), calculated based on the difference in exploration time between the incongruent object (the object placed in the incorrect context at test) and the congruent object (the object that matched the context based on prior training), normalized by the total exploration time. A higher DI reflects stronger memory for the correct object-context pairing.

We found that when the two training sessions were separated by 15 or 60 minutes, memory for the first object-context association was significantly impaired, while memory for the second association was preserved (**Fig. 1c**). However, when the interval was extended to 300 minutes, memories for both associations were maintained (**Fig. 1c**). This protocol allowed us to assess RI and revealed that such interference selectively affected the earlier memory trace when the two experiences occurred in close temporal proximity.

The serotonin type 2A receptor (5-HT2AR) is highly expressed in the cortex and has been extensively implicated in the control of memory interference during the retrieval of recognition memories (Bekinschtein et al., 2013; Morici et al., 2018). Then, we tested the hypothesis that enhancing cortical serotonergic signaling could prevent the forgetting of the first object-context association, potentially by supporting memory consolidation.

Rats were trained on the OIC task, using a fixed 60-minute delay between the two learning episodes. Immediately after the second learning session, animals received an infusion of DOI (a selective 5-HT2A/2CR agonist) or vehicle (VEH) into the medial prefrontal cortex (mPFC). Memory tests were conducted either 3 or 24 hours after DOI infusion to assess the effects of DOI on short and long-term memory, respectively (**Fig. 2a-b**).

**Figure 2:**
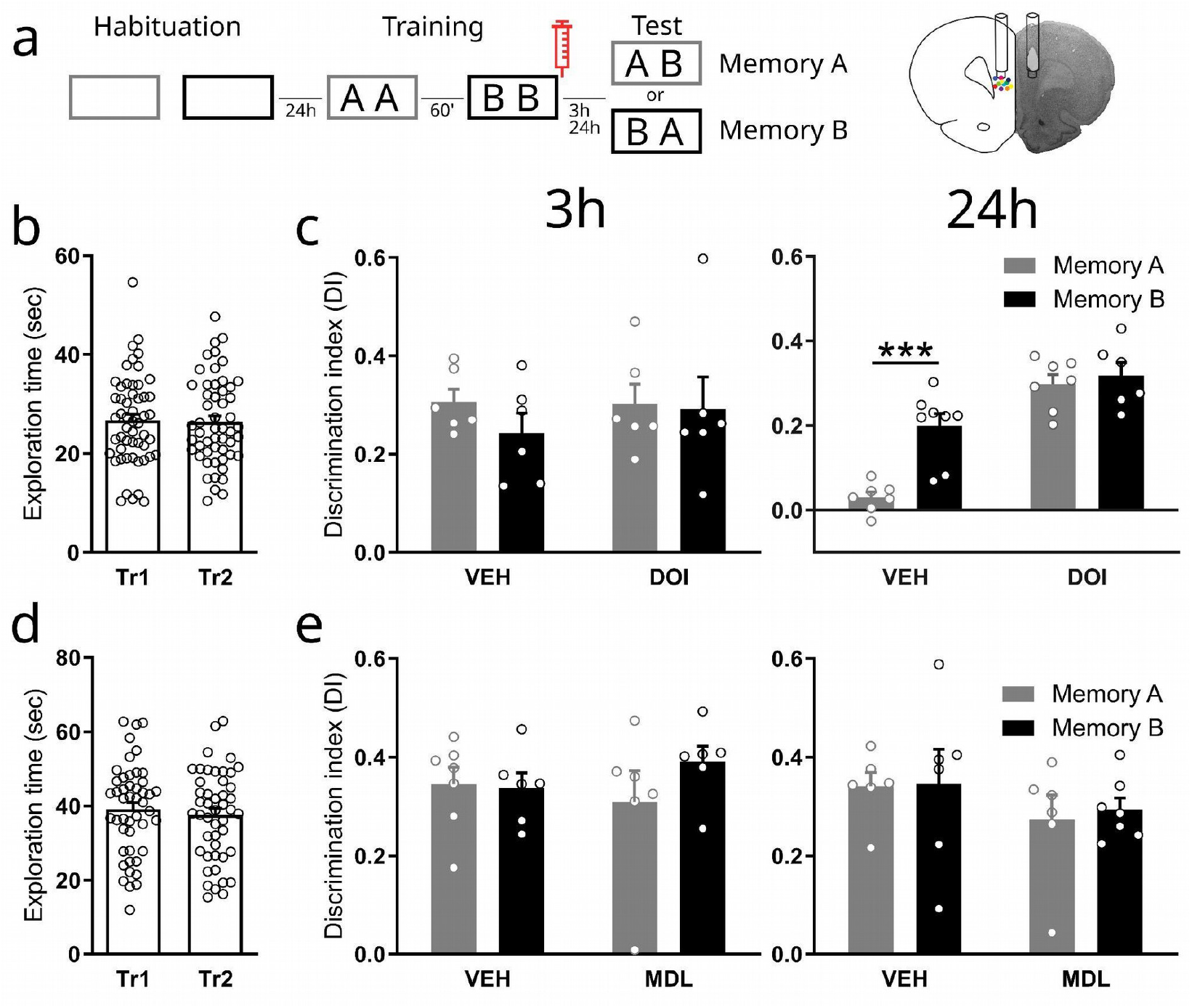
Activation of 5-HT2AR prevents the long-term dependent RI observed in OIC tasks. **(a)** Schematic representation of the infusion and behavioral paradigm. A selective 5-HT2AR agonist (DOI), antagonist (MDL), or vehicle (VEH) was delivered into the prefrontal cortex (PFC) immediately following the second acquisition session. Right panel: Coronal section of the cortical region we targeted for our pharmacological interventions. The weak or strong training protocols were used when DOI or MDL were employed, respectively. Short-term or long-term object-in-context (OIC) memories were tested 3 hours or 24 hours after the second learning session. (c, e) Exploratory time during each acquisition session. Paired t-test, p_b_=0.66, t_b_=0.44, p_d_=0.16, t_d_=1.44. (d, f) Discrimination Index (DI) obtained during the test sessions. White bars represent memory of the first learning (Memory A), while black bars represent memory of the second learning (Memory B). 2-way ANOVA followed by Sidak post hoc test. DOI experiment; 3h-delayed test, p_interaction_=0.57, F(1,20)=0.33; 24h delayed test, **p_interaction_=0.006, F(1,24)=8.88. ***p_post-hoc_<0,001. MDL experiment; 3h-delayed test, p_interaction_=0.27, F(1,21)=1.14; 24h-delayed test, p_interaction_=0.87, F(1,21)=0.02. n=6-8 per group.

Animals did not show differences in the total exploration time in either of the training sessions (**Fig. 2b**). We found that short-term memory was unaffected in both the VEH and DOI groups (**Fig. 2c, left panel**). In contrast, when we evaluated the response 24hs later, we found that while the VEH group exhibited the expected retroactive interference—demonstrating impaired memory for the first object-context pairing, the DOI group showed a recovery of that memory (**Fig. 2c, right panel**). These results suggest that retroactive interference primarily disrupted long-term memory consolidation of the first association. The 24-hour effect cannot be attributed to a general enhancement in memory consolidation, since DOI infusion after a brief 3-minute object–context exposure did not result in better performance than that observed in the VEH group at test. (VEH: 0.33 ± 0.04, DOI: 0.36∓ 0.03, unpaired t-test, p=0.58, t=0.57, n=5 per group). This suggests that 5-HT2AR activation in the mPFC can protect newly formed memories from retroactive interference, potentially by enhancing cortical consolidation mechanisms.

Since cortical 5-HT2AR activation was sufficient to prevent retroactive interference-induced forgetting, we next asked whether blocking 5-HT2AR signaling could, conversely, promote it. To test this, we used an extended training protocol previously validated as not inducing retroactive interference during consolidation (Bekinschtein et al., 2013; Morici et al., 2018). Immediately after the second training session, animals received an infusion of MDL 11,939 (a selective 5-HT2AR antagonist, MDL) or vehicle (VEH) into the medial prefrontal cortex.

Each object-context association was then tested for short- and long-term memory retention. Total object exploration times during training were comparable across groups and sessions (**Fig. 2d**), indicating no differences in general exploratory behavior. Furthermore, we observed no evidence of retroactive interference in either the short- or long-term memory tests (**Fig. 2e**). These results suggest that 5-HT2AR signaling is not required to maintain normal memory integration under strong encoding conditions and that its blockade is not sufficient to induce interference when memories are robustly formed.

Since DOI shows selectivity for both 5-HT2A and 5-HT2C receptors, we next wanted to evaluate if 5-HT2A receptors mediated the blockade of the RI process. To do so, we used a within-subjects design and trained rats in the ST version of the OIC paradigm while TCB-2 -a selective 5-HT2AR agonist-was infused in the mPFC either alone or co-infused with MDL. The animals were randomly assigned to one of the four experimental conditions (TCB-2, MDL, TCB-2 + MDL, VEH) and, in different weeks, each object-context association was evaluated. Each week, rats underwent a training session, and immediately after the second learning session, they received an infusion of TCB-2, MDL, TCB-2 + MDL, or VEH into the mPFC. The following day, memory for either the first or second object-context association was tested, with the tested association chosen pseudo-randomly across animals. One week later, the same procedure was repeated with new contexts and objects, using the same pharmacological condition assigned during the previous week. This time, the association that had not been tested the week before was tested.

We found that while the VEH and MDL groups exhibited the expected RI, this deficit was reversed when animals received TCB-2 infusions into the medial prefrontal cortex. Interestingly, this restored ability to recognize object-context association was lost in animals that received TCB-2 + MDL infusions (**Fig. 3c**). When evaluating the exploration times during training sessions, we observed that animals showed differences in the total exploration time in training sessions of the first week (**Fig. 3b**). However, we did not find differences when we analyzed exploratory times by treatment (2-way RM ANOVA. p_interaction_=0,9420, F(9,60)=0,3763), and more importantly no significant differences in TCB-2 group the one where RI is rescued (Tr1 vs Tr2, 1st week: p_post-hoc_=0,6853; 2^nd^ week: p_post-hoc_=0,9787), suggesting that the effect observed during the test session could not be explained by exploratory differences during acquisition.

**Figure 3:**
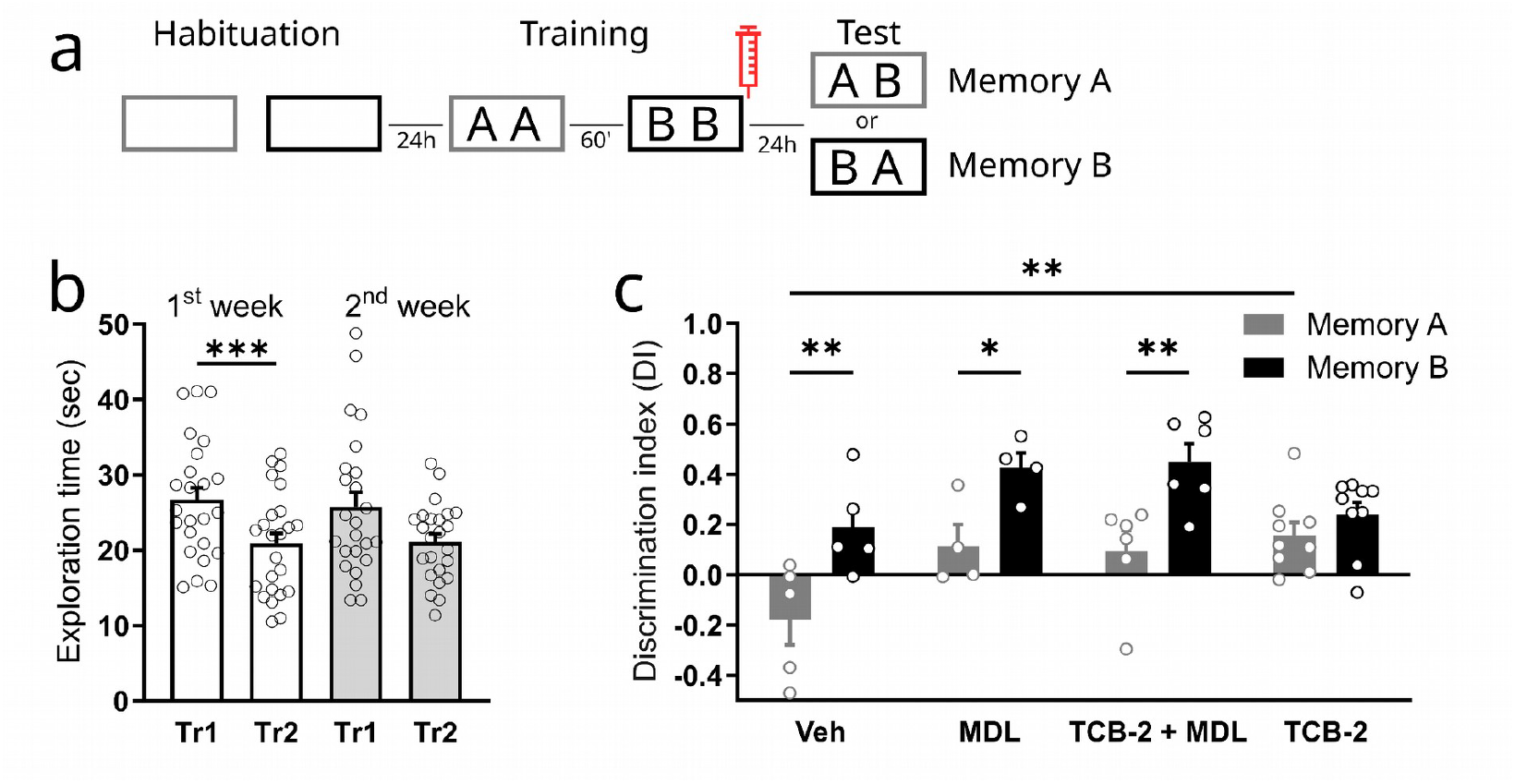
Selective activation of 5-HT2AR prevents the long-term dependent RI observed in OIC tasks. (a) Schematic representation of the infusion and behavioral paradigm. A selective 5-HT2AR agonist (TCB-2), antagonist (MDL), or a combination of both (TCB-2 + MDL), or vehicle (VEH) was delivered into the medial prefrontal cortex (mPFC) immediately following the second acquisition session. Long-term object-in-context (OIC) memories were tested 24 hours after the second learning session. (b) Exploration time for each object obtained during acquisition sessions. 2-way ANOVA followed by Holm-Sidak post hoc test ***p<0.0001; 1^st^ week Tr1 vs. 1^st^ week Tr2, ***ppost-hoc=0,0008. (c) Discrimination Index (DI) obtained during the test sessions. Gray bars represent memory of the first learning (Memory A), while black bars represent memory of the second learned object-context pair (Memory B).2-way RM ANOVA followed by Šídák’s post hoc test. Ctx, ***p<0,0001, F(1,20)=29,95; Drug, **p=0,0099, F(3,20)=4,956; p_interaction_=0,0974, F(3,20)=2,407. For Ctx factor (A vs. B): Veh, **p_post-hoc_=0,0026; MDL, *p_post-hoc_=0,0164; TCB-2+MDL, **_ppost-hoc_=0,0016; TCB-2, p_post-hoc_=0,3287. For Drug factor at Ctx A: Veh vs. TCB-2, **p_post-hoc_=0,0076. n=4-9 per group.

The test session in the object-in-context task assesses memory retention for an object-context association, as animals must detect that one of the objects is contextually incongruent. Because successful performance relies on the integration of both object and contextual information, we next asked whether retroactive interference disrupts not only the object-context association as a whole but also the memory of the object itself.

To test this, animals were trained using the weak OIC protocol. However, during the long-term memory test, instead of presenting a contextually incongruent object, we introduced a novel object (**Fig. 4a**). In this condition, the contextual component becomes less relevant, and the task is solved by recognizing the previously encountered object.

**Figure 4:**
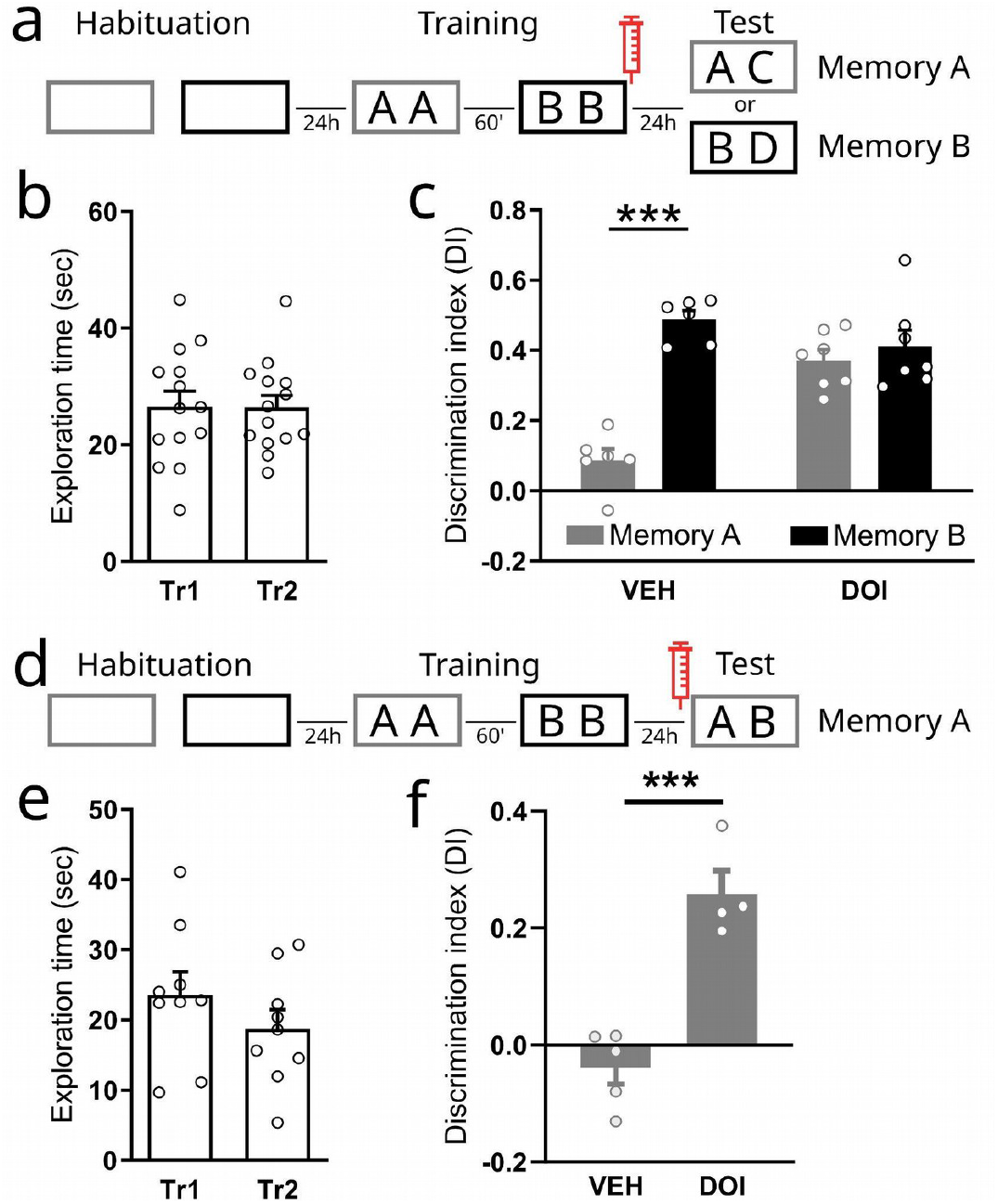
Time-Sensitive Modulation of Retroactive Interference by PFC 5-HT2AR Activation. (a) Schematic representation of infusions and behavioral paradigms. Animals were trained in the OIC task using the short-training protocol. Immediately after training, they received intracerebral drug infusions, and memory for one of the two original objects (A or B) was tested 24 hours later. (b) Exploratory time was obtained during acquisition sessions. Paired t-test, p=0.93, t=0.09. (c) Discrimination Index (DI) obtained during test sessions. White bars (A) represent the memory of Tr1, while violet bars (B) represent the memory of Tr2. 2-way ANOVA followed by Sidak post-hoc test. ****p_interaction_<0.0001, F(1,11)=32.17. ****p_post-hoc_ < 0,0001. N = 6-7 per group. (d) Schematic representation of infusions and behavioral paradigms. Animals were also trained on the OIC task using short training, and before the Test session, VEH or DOI was infused into the PFC to assess memory retrieval. (e) Exploratory time was obtained during acquisition sessions. Paired t-test, p=0.19, t=1.43. (f) Discrimination Index (DI) obtained during test sessions. Unpaired t-test, p=0.0004, t=6.20, N=4-5 per group.

Under basal conditions, animals failed to discriminate between the familiar object from the novel one, indicating impaired object memory from the first learning episode. Strikingly, this deficit was reversed when animals received DOI infusions into the medial prefrontal cortex immediately after training (**Fig. 4b-c**). These results suggest that retroactive interference affects the object memory from the first learning, and that serotonergic activation in the prefrontal cortex after training can rescue it.

These results indicate that RI occurs at least during the consolidation process. However, our results do not address whether the effects observed were due to a complete loss of memory or to a change in the accessibility of the first memory trace. To test this last hypothesis, we performed an experiment in which, in this case we infused DOI before the test session (**Fig. 4d**). During the Training session, all groups showed similar levels of object exploration (**Fig. 4e**). At Test, the VEH group exhibited RI-induced impairment in the expression of Memory A. Remarkably, activation of prefrontal 5-HT2AR during the Test session was sufficient to rescue the expression of this memory (**Fig. 4f**). This suggests that the RI-induced memory impairment does not result from disrupted consolidation. Instead, it reflects a failure to access Memory A at retrieval.

## Discussion

In this study, we investigated how serotonergic modulation through the 5-HT2A receptors in the prefrontal cortex influences memory when two competing experiences occur in close temporal proximity. Using the object-in-context (OIC) paradigm, we found that retroactive interference (RI) disrupts memory for a previously learned object-context association when a second experience occurs shortly after, if the training session was short. Strikingly, infusing a selective 5-HT2A receptor agonist into the medial prefrontal cortex immediately after the second learning experience was sufficient to prevent this interference-induced forgetting. Moreover, our findings reveal that RI affects at least the object-specific component of the memory trace.

Retroactive interference was initially defined as the impairment that occurs when a piece of information or a task is introduced between the presentation of target information and the subsequent recall of that information ^16–19^. The time-dependent stabilization of newly acquired memory, the memory trace, is vulnerable to disruption by a variety of amnestic influence or learning of new associations ^1^. Despite RI having been studied for decades, the circuits, pathways, and biochemical pathways involved are not completely understood. In rodents, RI has been reported, among other tasks, in object recognition paradigms ^3,4^, showing that the PFC plays a role in selection and interference resolution ^3^. In this work, by manipulating 5-HT2AR signaling, we have revealed a previously unreported role for serotonin in this process. We show that activating 5-HT2A receptors in the PFC immediately after the acquisition phase of the second association can similarly restore access to the original memory, reinforcing the idea that serotonergic signaling contributes, by modulating mPFC activity, in the stabilization of memories ^20–22^ under conditions of interference. Contrary, when the putative strength of the memories formed is high, 5-HT2AR activity does not appear to be necessary. These differences in 5-HT2AR requirement might suggest that triggering 5-HT2AR signalling might boost cellular pathways that might be needed for engrams differentiation. Also, it suggests the serotonergic system as an interesting pharmacological target for interference blockers. What’s particularly compelling is that this effect was evident not only when the receptor was engaged after the second learning, but also when it was activated just before memory retrieval. This points to two temporally distinct roles. for 5-HT2ARs. On the one hand, post-encoding activation may enhance synaptic plasticity processes, such as long-term potentiation, supporting the consolidation of otherwise weak memories ^23^. On the other hand, activation before recall appears to improve the precision of memory retrieval, likely by enhancing the signal-to-noise ratio in the underlying cortical networks ^24^. Interestingly, a recent study found that RI-induced amnesia could be reversed by optogenetically activating neuronal assemblies associated with the context of the first experience ^5^. This finding suggests that memory impairment following RI may be due to a retrieval deficit, consistent with the idea that pre-recall modulation enhances memory accessibility.

Taken together, these observations suggest that the strength of an experience could regulate 5-HT2AR expression or responsiveness within engram-relevant neuronal populations. Pharmacologically enhancing 5-HT2AR signaling, then, might serve dual functions: reinforcing the storage of vulnerable memories during consolidation, and improving their accessibility at the time of retrieval. Beyond its mechanistic implications, this dual role also highlights the receptor’s therapeutic potential for addressing memory disruption, particularly in conditions where interference or competition between experiences compromises memory fidelity.

## Acknowledgements

We thank Dr. Azul Silva for helpful discussions on the manuscript, Jaime David, and Vet. Maria Ines Bensansón for technical assistance.

## Author contributions

JFM, PB and NW designed the experiments, JFM, MI, PK, MBZ, and AS collected the data, JFM and NW wrote the manuscript. All authors reviewed the manuscript and approved the conclusions.

